# Neural harmonics of syntactic structure

**DOI:** 10.1101/2020.04.08.031575

**Authors:** Alessandro Tavano, Stefan Blohm, Christine A. Knoop, R Muralikrishnan, Lauren Fink, Mathias Scharinger, Valentin Wagner, Dominik Thiele, Oded Ghitza, Nai Ding, Winfried Menninghaus, David Poeppel

**Affiliations:** Department of Neuroscience, Max Plank Institute for Empirical Aesthetics, 60322 - Frankfurt am Main, Germany; Department of Language and Literature, Max Plank Institute for Empirical Aesthetics, 60322 - Frankfurt am Main, Germany; Department of Music, Max Plank Institute for Empirical Aesthetics, 60322 - Frankfurt am Main, Germany; University of Marburg, 35037 – Marburg, Germany; Department of Biomedical Engineering, Boston University, 02215 - Boston, USA; College of Biomedical Engineering and Instrument Sciences, Zhejiang University, 310027 - Hangzhou, Zhejiang Province, China; Department of Psychology, New York University, 10003 - New York City, New York, USA; Enrst Strüngmann Institute, Frankfurt am Main, Germany

## Abstract

Can neural rhythms reflect purely internal syntactic processes in multi-word constructions? To test this controversial conjecture - relevant to language in particular and cognition more broadly - we recorded electroencephalographic and behavioural data as participants listened to isochronously presented sentences of varying in syntactic complexity. Each trial comprised ten concatenated sentences and was either fully grammatical (regular) or rendered ungrammatical via randomly distributed word order violations. We found that attending the regular repetition of abstract syntactic categories (phrases and sentences) generates neural rhythms whose harmonics are mathematically independent of word rate. This permits to clearly separate endogenous syntactic rhythms from exogenous speech rhythms. We demonstrate that endogenous but not exogenous rhythms predict participants’ grammaticality judgements, and allow for the neural decoding of regular vs. irregular trials. Neural harmonic series constitute a new form of behaviourally relevant evidence for syntactic competence.

## Introduction

Speech and language are related but distinct dimensions of verbal communication [1—4]. In a nutshell, speech is linear, while language is hierarchically organized. Temporal regularities in speech may be relevant for language production and comprehension [3, 5—13]): for example, entrainment to syllabic rhythm may contribute to word perception [3,6,14]. However, the core of language processing lies in the tacit, internal knowledge of syntactic rules, which combine words into meaningful utterances [15]. The challenge is how to infer hierarchy from the sequence of neural responses to speech units. While valuable information has been obtained by measuring event-related potentials responses (ERPs) to syntactic rule violation [16–17], ERPs reflect a mixture of functional and physical properties of individual words and syntactic structures [18–19], making it difficult to capture the unfolding of syntactic competence over multiple words. To go the extra mile, a link can be established between the duration of multi-word syntactic units, whose extraction presupposes competence, and slow brain rhythms. Ding and colleagues [20] made such a connection by asking native speakers of English and Mandarin Chinese to listen to trials of ten spoken sentences composed of monosyllabic words presented at a fixed rate of 4 Hz. Forcing word isochrony into sentence perception allows to ‘frequency tag’ [21] the duration of abstract syntactic units which regularly repeat albeit with ever differing words. Ding and colleagues [20] tested only one syntactic structure: Two-word noun phrases (NP = a phrase headed by a noun, such as “My nose”) followed by two-word verb phrases (VP = a phrase headed by a verb, such as “is big”, see Figure 1a). A Fast Fourier Transform (FFT) analysis highlighted significant peaks of spectral energy not only at 4 Hz, that is word frequency, but also at 2 Hz and 1 Hz, which the authors interpreted as independently tracking the period of phrases (2-word units = 2 Hz) and sentences (4-word units = 1 Hz). Of note, linguistic rhythms emerged only if participants listened to their own language [22]. Does the hierarchy of neural rhythms obtained by Ding and colleagues [20] reflect the online incremental combination of words into phrases, and phrases into sentences [23]? If this held true, then it would define a direct mapping between neural rhythms and levels of syntactic structure, bypassing the problem of linearization.

**Figure 1.**
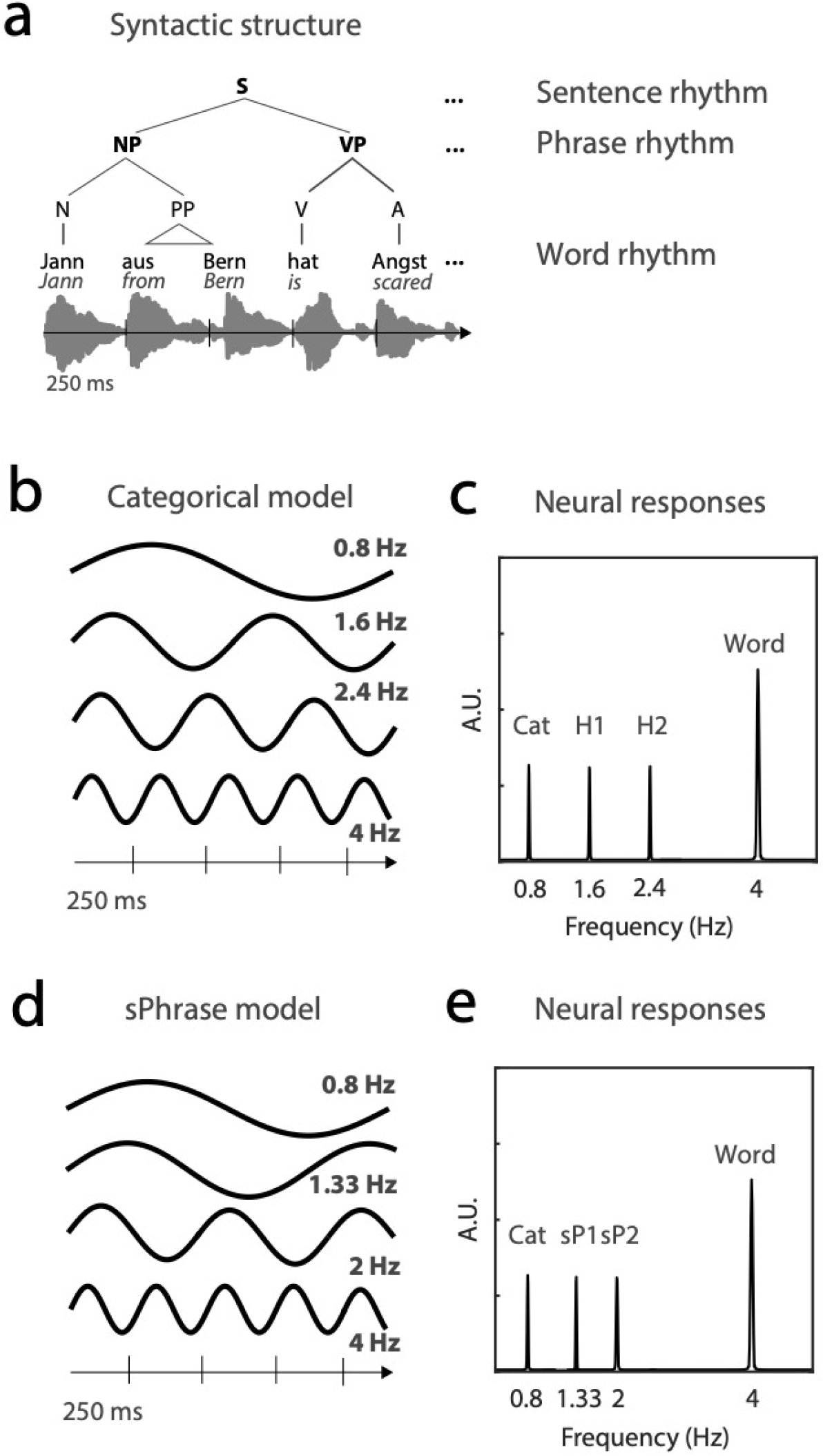
Stimulus structure and neural response models: a. A syntactic description of a sample sentence structure which, when presented in chains of ten sentences with the same structure but realized using different lexical items, would permit distinguishing harmonic peaks from sPhrase peaks in the frequency domain. Three levels of linguistic investigation are highlighted: word (stimulus rate), phrase and sentence. b. For five-word sentences running at 4 Hz, the categorical model assumes that NPs and VPs repeat every 0.8 seconds, on a pair with sentences. c. Consequently, an FFT analysis of neural data should highlight — beside a word rate peak — a 0.8 Hz peak of energy, representing the categorical rhythm (Cat), and its harmonics (first harmonic — H1 — 1.6 Hz, second harmonic — H2 — 2.4 Hz). d. The sPhrase model would suggest instead the existence of phrasal tracking blind to their lexical head, and therefore yielding, in the example at stake, a 2 Hz rhythm (for the two-word NP) and 1.33 Hz rhythm (for the three-word VP). e. The neural responses of sentences and phrases would be independent.

We argue that the hypothesis of an isomorphism between syntactic and neural rhythms, however attractive, is invalid. If the human brain tracks phrasal node types (e.g., NP, VP, etc. [24–29]), in a frequency tagging experiment each phrasal at a given sequential position repeats only once every sentence period. This means that individual phrasal nodes, as well as sentence nodes, contribute to the same rhythm, generated by the sentence period, which we term *Categorical* rhythm, as it reflects the repetition of abstract syntactic categories. If this held true, it would be impossible for phrases to generate their own, independent rhythm. Hence, previously defined “phrasal rhythms” [20, 22–23] would simply reflect a first harmonic process of the categorical rhythm (see Figures 1a,b,c), effectively barring any direct, one-to-one mapping between linguistic and neural hierarchies.

An independent phrasal rhythm could emerge if and only if there exited a superordinate phrasal category, which we term *sPhrase*, which would cycle through all phrase type, blind to their lexical head type (that is, VP and NP would not be distinguished: see Figures 1d,e). The idea that phrase structure is independent of lexical categories has exerted a large influence in mainstream syntactic theory [30], and although rebutted [31], it has never been tested on neural responses.

We pitched Categorical and sPhrase models against each other by systematically varying the size of phrasal units in six sentence structures in the German language. We recorded encephalographic (EEG) and behavioural data while 31 participants listened to trials composed of ten isochronized sentences, and discriminated between regular trials (50%), which contained only grammatical sentences, and irregular trials, in which two successive sentences were rendered ungrammatical by violating word order (random word distribution).

Results lend no support to the existence of a superordinate sPhrase category. Attentively listening to regularly repeated abstract syntactic categories generates a Categorical rhythm with its own harmonic structure. Hence, brain rhythms are not isomorphic to syntactic units. However, the harmonics of neural categorical rhythms are mathematically independent from word rate, for the first time we can dissociate bottom-up entrainment to word rate from top-down syntactic rhythms. The amplitude of neural responses to categorical harmonics significantly predicts participants grammaticality judgements, and — in contrast to word rate — successfully classifies grammatically regular from irregular trials.

## Results

### Grammatical judgments

Trials (ten concatenated sentences) were organized into three contexts: 1) fast four-word sentences, with monosyllabic words running at 4 Hz; 2) fast five-word sentences, with monosyllabic words running at 4 Hz; 3) slow five-word sentences, built from disyllabic words running at 2 Hz. Within each context, two sentence structures were created by varying the relative duration of noun and verb phrases, in order to test their import on neural responses (see Figure 2).

**Figure 2.**
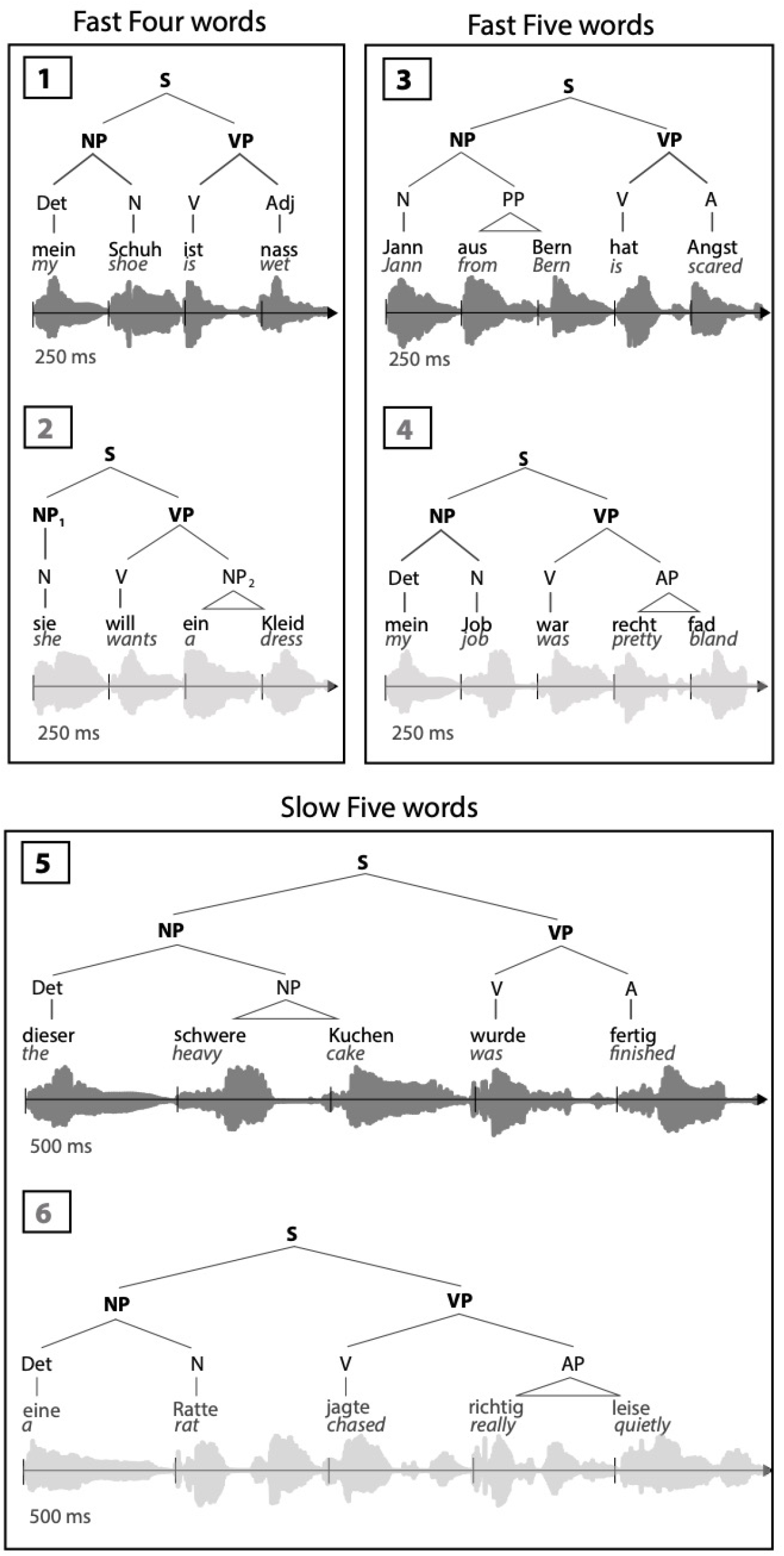
Sentence contexts: Sentence structures 1 and 2 have four monosyllabic words running at a 4 Hz, with structure 1 composed of a two-word NP and a two-word VP - replicating the stimuli of Ding et al. [20] -, and structure 2 of one-word NP followed by a three-word VP. Structures 3 and 4 have five monosyllabic words running at 4 Hz: Structure 3 sentences are composed of a three-word NP followed by a two-word VP, while structure 4 sentences flips that pattern. Structures 5 and 6 replicate structures 3 and 4 with disyllabic words running at 2 Hz. Syntactic trees of exemplary sentences and corresponding waveforms are provided for illustrative purposes. Black and grey colours identify even and odd sentence structures within each context (1-2, 3-4, 5-6, respectively). Notice that although current theories see determiners as heads of the determiner phrases (DP), with NPs as complements, for our purposes we follow a more traditional labelling convention. See Supporting Information for a complete list of sentences (N = 300).

Using a signal detection approach [32], we estimated task sensitivity and response bias for each sentence structure. The discriminability index (d’) measures task sensitivity: For all sentence structures, participants successfully distinguished irregular from regular trials, suggesting that the grammatical nature of the task was well understood: all ts_(30)_ ≥ 7.94, all ps ≤ 7.205*10^-09^, FDR-corrected (threshold p = 0.008. See Figure 3, left panel). There was a significant difference among sentence structures: F_(5,185)_ = 4.16, p = 0.001, η^2^ = 0.04. When FDR-corrected (threshold p = 0.006), type 1 differed from type 4, t_(30)_ = −3.34, p = 0.001, suggesting that sensitivity for structure 1 (mean = 1.83, SD = 1.24) was lower than for structure 4 (mean = 2.67, SD = 1.75). Response bias, measured by the decision criterion, showed that participants tended to deem irregular trials as regular for sentence types 1, 4, 5 and 6: all ts_(30)_ ≥ 3.65, all ps ≤ 9.749*10^-04^, FDR-corrected (threshold p = 0.021. See Figure 3, right panel). This suggests a conservative bias in grammaticality judgment. From this, we can infer that that the impoverished acoustic quality of stimuli following the pitch-flattening procedure did not confound participants’ comprehension. No significant difference in criterion estimates among sentence structures was found, suggesting that the stimulus generation procedure was bias-free: F_(5,185)_ = 1.81, p = 0.115.

**Figure 3.**
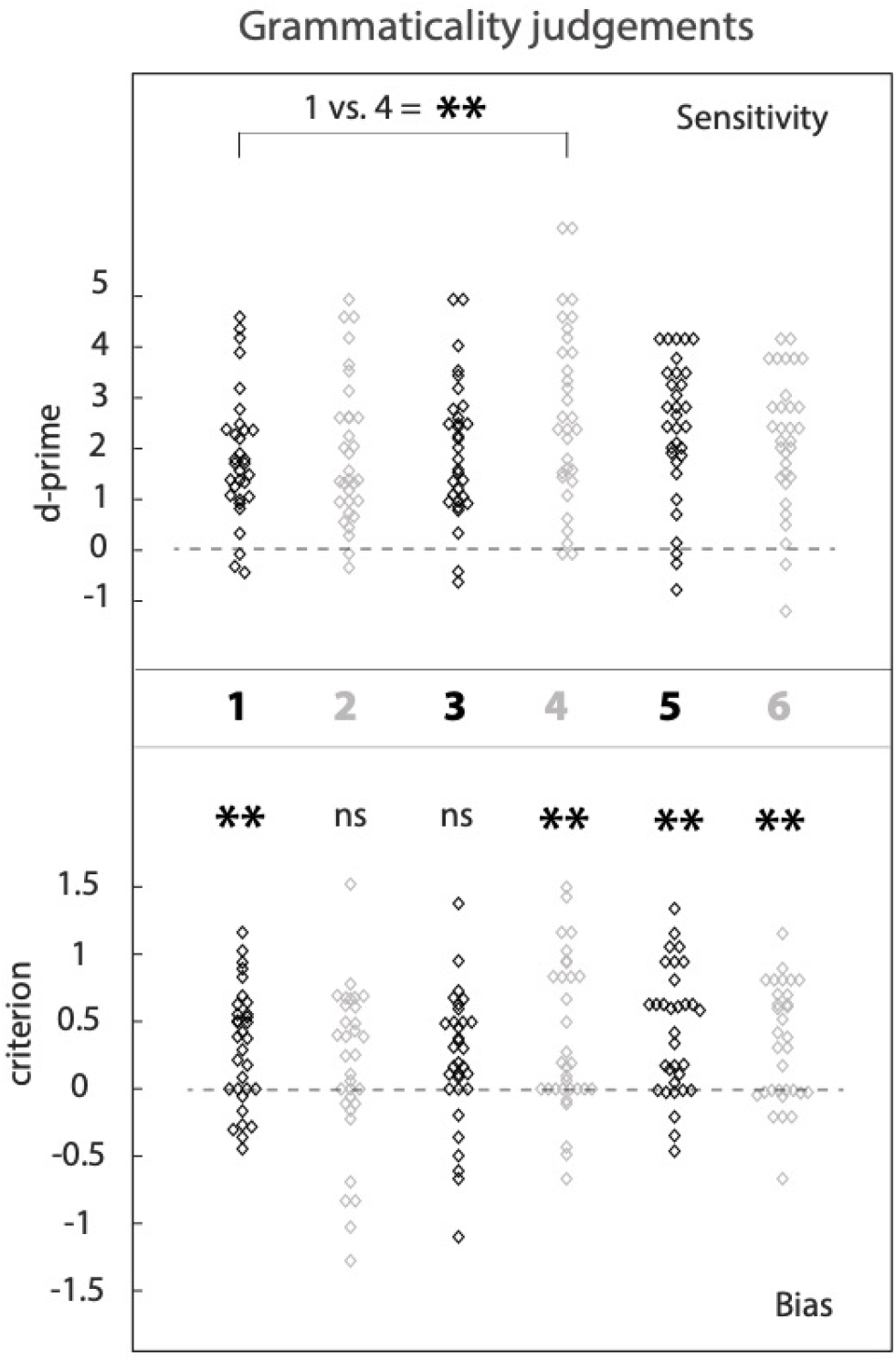
Behavioural accuracy. Upper panel: Participants discriminated between regular and irregular trials for all six sentence structures (middle bar, color-coded). A significant difference between structures 1 and 4 suggests that discriminating irregular from regular trials in structure 1, the original condition of Ding et al. [20], was more difficult than in structure 4. Middle panel: sentence structure label. Lower panel: A positive bias for structures 1, 4, 5 and 6 suggests that participants adopted a conservative criterion when operating grammatical judgments: They were more likely to respond that a given trial was regular, rather than irregular.

### Controlling for speech and acoustic cues

Phonemic transitions between successive words could implicitly cue phrasal segmentation. To verify whether this was the case, we calculated phonemic transition probabilities at critical boundaries within and between phrases in all experimental sentences, and found no significant differences (See Supporting Information, Figure S1a,b). Word frequency differed across sentence structures (Figure S1c), as expected due to the asymmetry in the distribution of monosyllabic and disyllabic words in natural languages (for the German language, see [33]).

Acoustic waveforms were subject to a Fast Fourier Transform analysis on a trial-by-trial basis. We extracted normalized power estimates using the complex modulus, and then calculated signal-to-noise ratio estimates (SNR). Results suggest the presence of a significant word rate effect for all sentence structures: all ts_(30)_ ≥ 55.04, all ps ≤ 1.080*10^-31^, FDR-corrected (threshold p = 0.001, Figure S2a,b,c). No other significant rhythmic components were found, suggesting that acoustic input could not drive higher-order, syntactic rhythms (see Supporting Information).

### Categorical harmonics

For neural data, we calculated inter-trial phase coherence estimates as the ratio of the FFT output and the amplitude of the signal, normalized by the number of trials for each sentence type and participant (See Materials and Methods). Neural frequency tagging effects were investigated using all trials, regardless of grammaticality. Figure 4, left column, illustrates phase coherence peaks for each sentence structure. Statistical analyses were run on standardized phase coherence signal-to-noise ration (SNR) scores. T-tests to zero showed robust entrainment to word rate for all sentence structures: all ts_(30)_ ≥ 9.21, all ps ≤ 1*10^-08^, FDR-corrected (threshold p = 0.008), and no significant difference between sentence structures within each context: all ts_(30)_ ≤ 0.27, all ps ≥ 0.788. Similarly, a significant harmonic series, here defined by the emergence of a categorical rhythm and its first two harmonics, H1 and H2, was found for all sentence structures: Category, all ts_(30)_ ≥ 3.61, all ps ≤ 0.001; H1, all ts_(30)_ ≥ 3.28, all ps ≤ 0.002; H2, all ts_(30)_ ≥ 3.09, all ps ≤ 0.004 (all FDR-corrected, threshold p = 0.008; See Figure 4, middle column). Hence, differences in the internal phrasal organization of sentences did not influence the presence/absence of a harmonic series. Crucially, the duration of abstract categories in sentence structures 3, 4, 5 and 6 ensures that first and second harmonic processes cannot be mathematically derived as subharmonics of stimulus rate [34]: for example, for sentence types 3 and 4 there exists no integer factor which multiplied by the first harmonic at 1.6 Hz would yield the 4 Hz word rate. The same holds for sentence types 5 and 6. This simple fact grants functional independence of each harmonic series relative to exogenous word rhythm, and proves the existence of purely endogenous, top-down rhythms.

**Figure 4.**
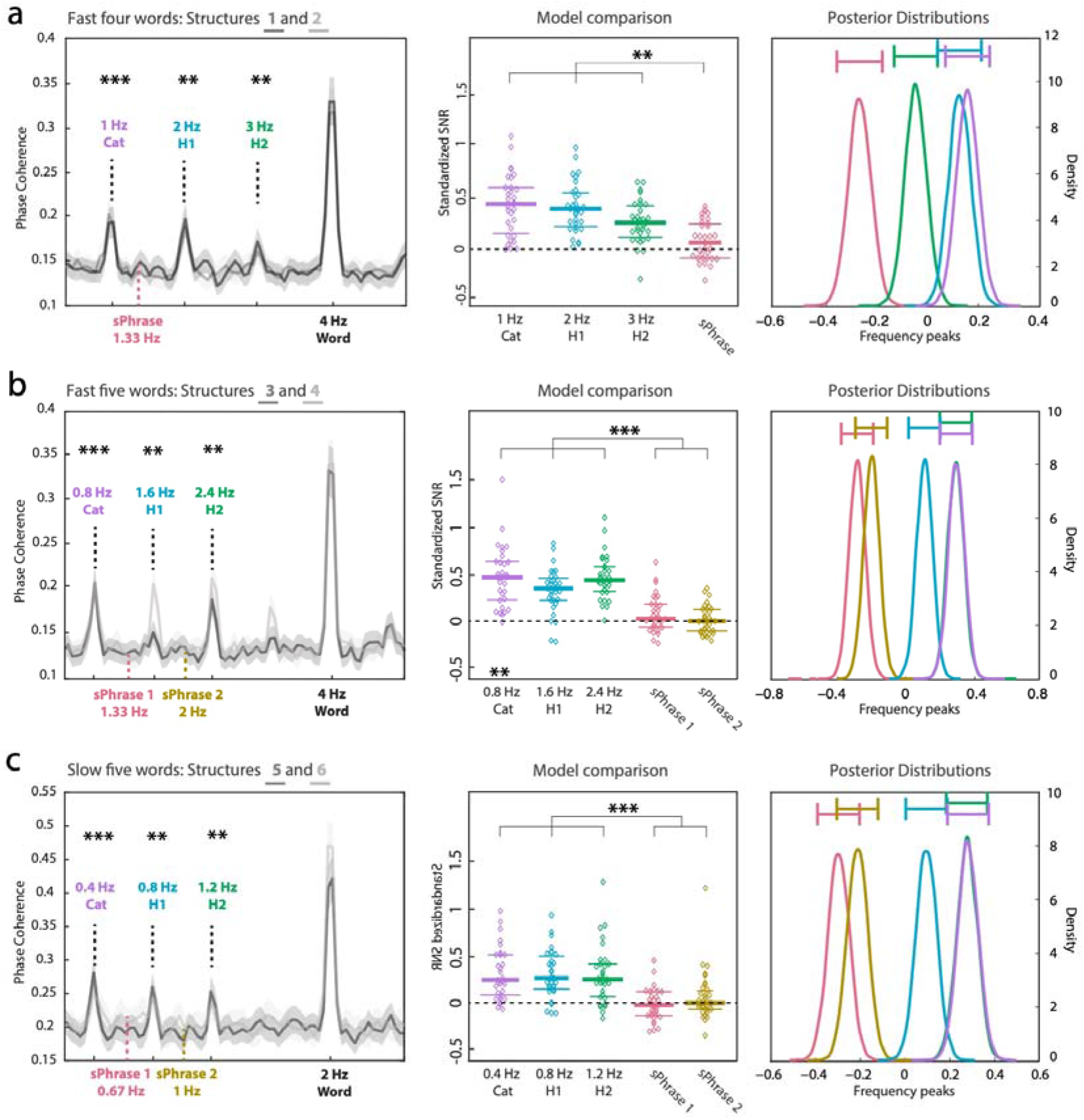
Neural rhythms. Left column shows phase coherence results for all trials, for each sentence context (sentence structure pairs: 1-2, 3-4, 5-6). Categorical rhythms and their harmonics, along with the positional distribution of possible *sPhrase* rhythms, are color-coded and indicated by vertical hyphenated bars. No significant peaks were found for any of the hypothesized sPhrase rhythms. Conversely, evidence for a categorical harmonics model was found for all sentence structures. The central column confirms the difference between categorical and sPhrase frequencies in all sentence contexts, using standardised SNR estimates. All probability values, here and in the text, are deemed significant or not within an FDR approach. Significance is marked f as follows: *** ≤ 0.0001, ** ≤ 0.001, ns = non-significant. Horizontal bars on right column figures represent 95% posterior odds credible intervals.

Syntactic variability within each sentence context modulated the strength of harmonic peaks. For the fast four-word context, a main effect of peak strength was found: F_(2,185)_ = 4.83, p = 0.01, η^2^ = 0.04. H2 differed from both categorical rhythm and H1: all ts_(30)_ ≥ 2.75, all ps ≤ 0.01. The responses for categorical rhythm (mean = 0.47 standardized SNR, SD = 0.35) and H1 (mean = 0.44, SD = 0.29) were larger than for H2 (mean = 0.27, SD = 0.25). In the fast five-word context, a significant peak by sentence structure interaction (F_(2,185)_ = 5.06, p = 0.009, η^2^ = 0.03) highlighted a difference in H1 responses between sentence structures 3 and 4: t_(30)_ = −4.08, p = 3.618*10^-05^. H1 in structure 3 (mean = 0.22, SD = 0.33) was smaller than in structure 4 (mean = 0.63, SD = 0.45). In the slow five-word context, a main effect of sentence structure was found: F_(1,185)_ = 14.16, p = 0.007, η^2^ = 0.05. Neural responses were larger in structure 5 (mean = 0.48, SD = 0.07) than in structure 6 (mean = 0.26, SD = 0.04).

### Model comparison

Sentence structure 1 is based on the original stimuli of Ding et al. [20]. Consequently, sPhrase and H1 coincide, making sentence structure 1 uninformative in adjudicating between categorical harmonics and sPhrase models. In sentence structure 2, the sPhrase model would expect a 1.33 Hz neural rhythm stemming from 3-word VPs (the 4 Hz rhythm stemming from the one-word NP is confounded with the stimulation frequency; Figure 4a, left column). Structures 3 to 6 allow for a full test of the internal generation of neural rhythms, as each phrase should create its own sPhrase peak (Figures 4b and 4c, left column; See also Figure 1).

T-tests to zero showed that no sPhrase peak reached significance: all ts_(30)_ ≤ 0.71, all ps ≤ 0.062, We then averaged SNR values across sentence structures (Figure 4, middle column), and run a series of one-way rmANOVAs, one for each sentence context, with peak frequency as a fixed factor. In all cases, categorical harmonic rhythms differed from sPhrase rhythms: all Fs_(3,123)_ ≥ 6.76, all ps ≤ 0.001, all η^2^ ≥ 0.13. Post-hoc comparisons showed that categorical harmonic structures outperformed sPhrase rhythms: all ts_(30)_ ≥ 2.73, all ps ≤ 0.01, FDR-corrected (threshold ≥ 0.016).

We evaluated the amount of evidence provided by phase coherence data using a Bayesian rmANOVA. The Bayes factor indicates that the data - across sentence contexts - were 156.786 times more likely under the model that includes peak frequency as a fixed factor, compared to the null model, which assumes no difference between peaks. Post-hoc comparisons highlight the difference between sPhrase and categorical harmonic peaks in the distribution of model-averaged posterior odds (normalized to the mean of all data). The sPhrase peak vs. H2 in the fast, four-word context revealed posterior odds of 1.782 against the null hypothesis, which indicates moderate evidence in favour of the alternative hypothesis (BF_10_ = 4.301). For all remaining comparisons in the fast four-word context, as well as in the remaining sentence contexts, the evidence in favour of the alternative hypothesis was very strong: all BF_10_ ≥ 47.309 (Figure 4, right column).

### Grammaticality in the frequency domain

We have determined that the brain encodes the duration of regularly repeated abstract syntactic nodes – NP, VP, sentence – within a harmonic structure of their period. However, the emergence of significant peaks of neural activity does not imply that they encode perceived grammaticality. To test this, we re-calculated SNR estimates of inter-trial phase coherence separately for hits (correctly identified irregular trials) and correct rejections (correctly identified regular trials). Omnibus rmANOVAs with factors frequency peak, sentence structure and grammaticality, separately for each sentence context, highlighted a significant grammaticality effect: all Fs_(1,371)_ ≥ 9.50, p ≤ 0.01, η^2^ = 0.02. In all cases, phase coherence for grammatical trials was larger than for trials containing ungrammatical sentences: fast four words, grammatical mean = 0.23, SD = 0.16, ungrammatical mean = 0.11, SD = 0.16; fast five words, grammatical mean = 0.35, SD = 0.23, ungrammatical mean = 0.21, SD = 0.16; slow five words, grammatical mean = 0.26, SD = 0.14, ungrammatical mean = 0.24, SD = 0.13 (see Figure 5a). For the fast five words context, a significant difference between types was also found, mirroring model comparison results: F_(1,371)_ = 14.53, p = 0.0006, η^2^ = 0.04. No other significant main effect or interaction was found: all Fs_(1,371)_ ≤ 2.60, p ≥ 0.08.

**Figure 5.**
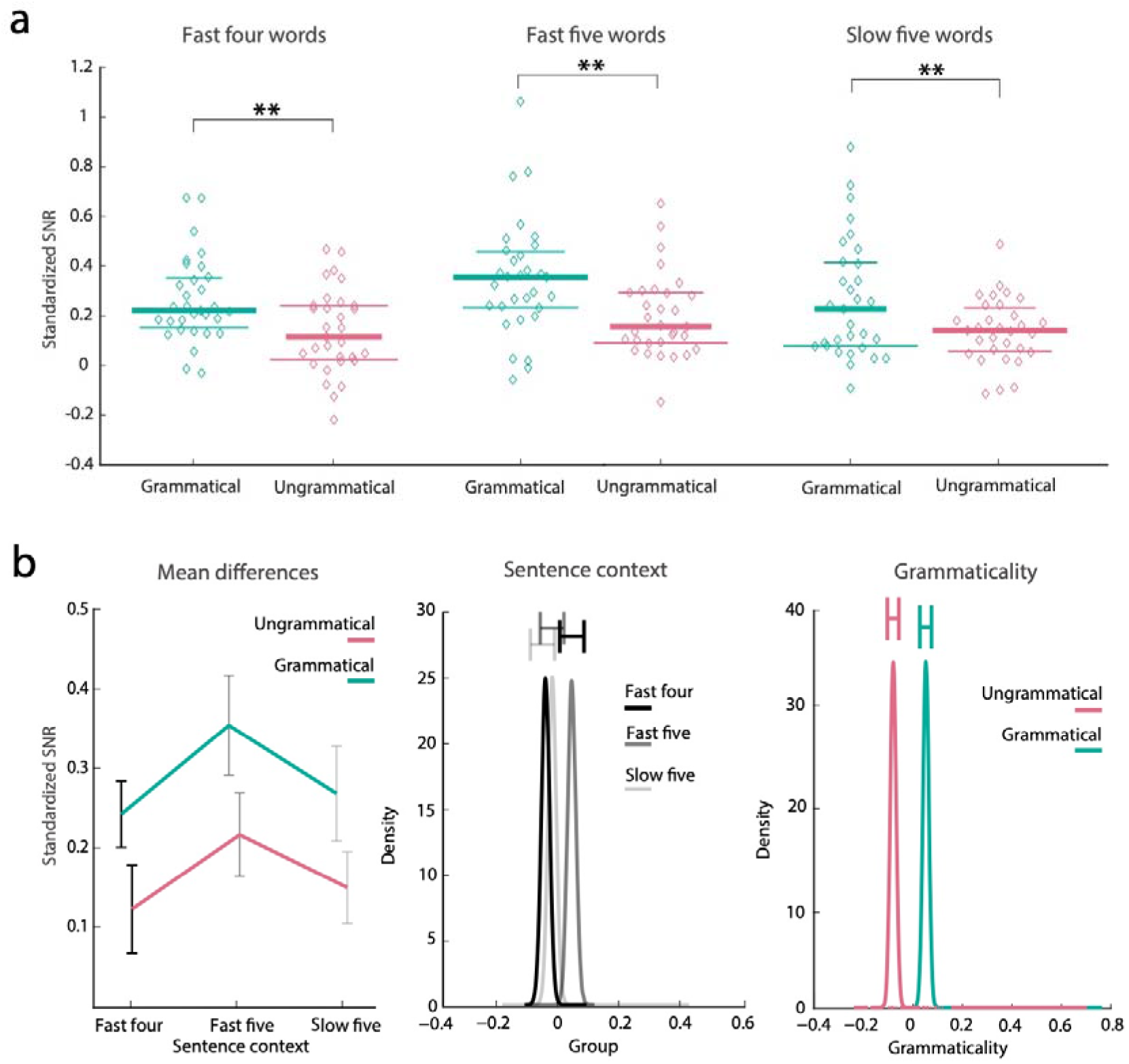
Neural rhythms of grammaticality: a. An omnibus rmANOVA analysis on averaged endogenous rhythms shows a significant effect of grammaticality in each sentence context. b. A Bayesian rmANOVA highlights two main effects: sentence context and grammaticality. The grammaticality effect provides decisive evidence for endogenously-driven phase coherence. The sentence context effect provides decisive evidence suggesting that the fast five-word context generates the largest phase coherence scores. Significance is marked as follows: *** ≤ 0.0001, ** ≤ 0.001, ns = non-significant. Horizontal bars on right column figures represent 95% posterior odds credible intervals.

Using a Bayesian rmANOVA with factors sentence context and grammaticality (Figure 5b, left panel), the Bayes factor indicates that the data were at least 1.185*10^07^ times more likely under the model that includes grammaticality and sentence context as main factors: updated probability = 0.89, prior null model probability = 0.20. The evidence for the interaction component was very small, as adding it changed prior odds only marginally - BF_M_ = 0.433 -, as compared to simply adding main effects: BF_M_ = 32.352. Post-hoc comparisons of grammatical vs. ungrammatical trials revealed a BF_10_ of 40847.319 against the null hypothesis, which indicates decisive evidence in favour of the alternative hypothesis. As for the context factor, decisive evidence was obtained for the difference between fast five- and four-word sentence contexts (BF_10_ = 394.811, uncorrected for multiple comparisons), and strong evidence for the difference between fast and slow five-word sentences (BF_10_ = 17.039, uncorrected). We conclude that the strength of categorical harmonic structures, resulting from attending to the repetition of abstract syntactic categories, is modulated by trial grammaticality.

### Harmonics encode grammaticality judgements

When participants decide whether a trial contained ungrammatical sentences or not (irregular trial), their judgment must be driven by implicit syntactic competence of the German language. However, violations of expected word order also represent a form of sensory surprise, which per se can capture attention [35] and independently predict behavioural accuracy. To partition these two contributions, we created two simple neural predictors: an endogenous one, by calculating for each participant the mean standardized phase coherence score across categorical harmonic peaks (Cat, H1 and H2), for all sentence structures, separately for correct rejections and hits; and an exogenous one, by averaging the strength to which each participant entrained to external word rate, again across sentence structures and separately for regular and irregular trials. Each neural predictor was then regressed onto task sensitivity scores, averaged across all sentence structures. A functionally relevant dissociation emerged between endogenous and exogenous neural rhythms extracted from regular vs. irregular trials. For fully grammatical, regular trials, the endogenous index significantly predicted participants’ ability to decide whether a trial was regular or irregular: F_(1,29)_ = 15.93, p = 4.096*10^-04^, adjR^2^ = 0.33. However, the exogenous index did not: F_(1,29)_ = 0.70, p = 0.407 (see Figure 6a). This suggests that only endogenous categorical harmonics reflect access to internal models of syntactic knowledge. For irregular trials, however, both indexes significantly predicted participants’ ability to decide whether a trial was regular or irregular: endogenous, F_(1,29)_ = 23.15, p = 4.277*10^-05^, adjR^2^ = 0.42; exogenous, F_(1,29)_ = 13.68, p = 0.0011, adjR^2^ = 0.28 (see Figure 6b). To decide whether the results for exogenous rhythms in irregular trials reflect sensory surprise or grammatical violation detection, we resorted to a neural decoding approach. We calculated standardized phase coherence scores separately for each relevant frequency peak – categorical, H1, H2 and word rate –, and used a linear discriminant analysis (LDA) with electrodes as a feature dimension to classify hits and correct rejection trials from neural rhythms. The performance of the classifier – scored by computing the nonparametric measure of the area under the curve (AUC) of the receiver-operating characteristic – was significant for all rhythms belonging to categorical harmonic series: Categorical rhythm = .58, CI: 0.47 – 0.53; H1 = .64, CI: 0.46 – 0.52; H2 = .65, CI: 0.46-0.53. Crucially, however, it was at chance for entrainment to word rate: .50, CI: 0.47-0.53 (see Figure 6c). Hence, we conclude that only top-down rhythms, constituting a harmonic series of syntactic node repetition period, encode information about a trial grammaticality.

**Figure 6.**
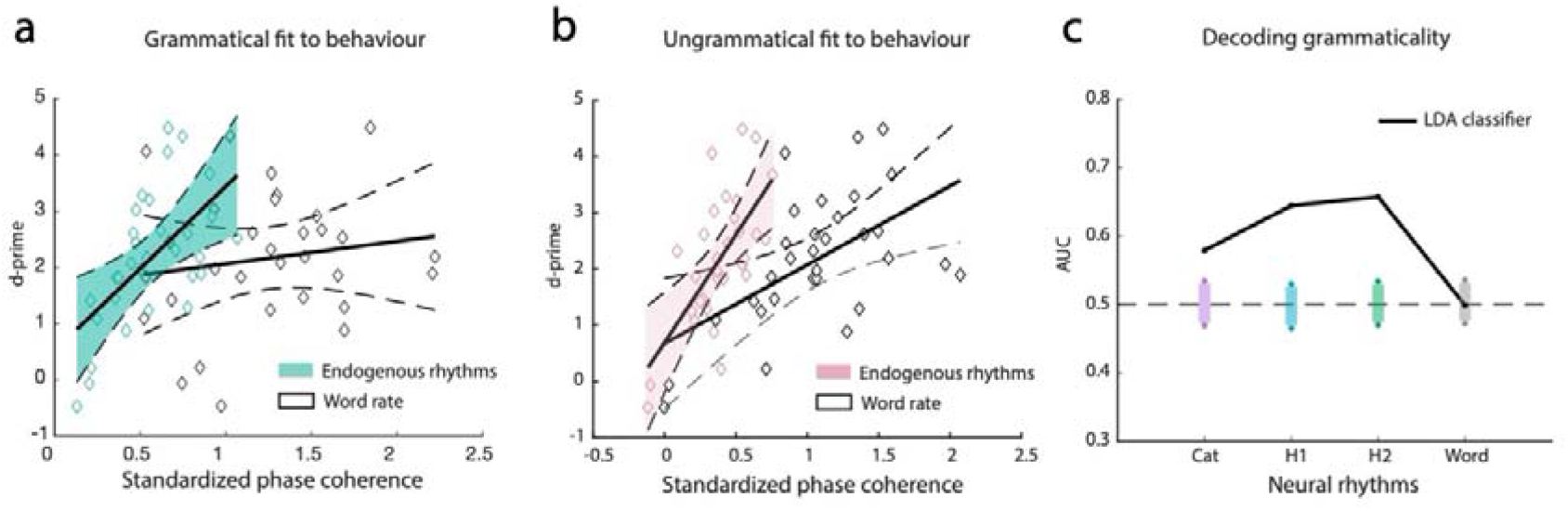
Brain/behaviour fit: a. Accuracy fit of endogenous and exogenous rhythms extracted from regular trials. Only endogenous rhythms predict behavioural accuracy. Thick lines represent best fit. Thin, dotted lines indicate each regression’s 95% confidence interval. b. When extracted from irregular trials, both endogenous and exogenous rhythms significantly predict grammaticality judgements, suggesting the contribution of both syntactic competence and sensory surprise to neural activity. c. However, AUC scores from an LDA analysis show that it is possible to correctly classify regular vs. irregular trials using endogenous rhythms, but not entrainment to word rate. Shaded color-coded areas report 95% confidence intervals, obtained via Montecarlo resampling. Significance: *** ≤ 0.001, * ≤ 0.05, ns = non-significant.

## Discussion

We aimed at extracting a neural signature of purely internal syntactic information from multi-word speech units. Since speech acoustic and linguistic information are intertwined in verbal communication, to zoom in on syntactic competence we reduced the perceptual significance of speech cues to comprehension by forcing words to onset at regular intervals. This experimental strategy may appear artificial, because intuitively we associate the naturalness of language to the naturalness of speech. However, once the difference between the hierarchical nature of syntax vs. the linear nature of speech is highlighted, it becomes evident that isochronizing speech leaves the core of language competence, and hence its naturalness, intact. That intuition founded Ding et al.’s [20] milestone attempt to go beyond single-word neural responses (e.g., ERPs), and measure neural rhythms to multi-word language units. We provide evidence that frequency tagging goes deep in that it captures internal syntactic processes relevant for grammaticality judgements. However, this result does not rest on an isomorphic mapping between the levels of syntactic analysis (word, phrase, sentence) and the hierarchy of neural rhythms (word rhythm, phrase rhythm, sentence rhythm). Frequency tagging does not allow to separately read out sentences and phrases, because their abstract syntactic nodes are repeated with the very same period (for a congruent lexical approach, see [36]). The repetition of linguistic categories - both phrases and sentences - with the same period generates a categorical neural rhythm, which gives rise to a harmonic series that encodes grammaticality judgements. Thus, harmonic series represent an indirect mapping between syntactic competence and neural rhythms.

Importantly, our experimental manipulations document that the harmonic series generated by category-driven syntactic rhythms are independent of entrainment to word rate. For example, in conditions 3 and 4, the first and second harmonics at 1.6 and 2.4 Hz cannot be mathematically derived as sub-harmonic processes of the 4 Hz word rate. This strongly supports a principled distinction between endogenous rhythms, which follow abstract node repetitions and encode grammaticality, and exogenous rhythms, which entrain to word rate and likely encode sensory surprise, when extracted from irregular trials (See Figure 6c).

Notice that categorical rhythms and their harmonics emerge regardless of internal sentence structure, although the latter modulates the strength of the harmonics, possibly highlighting differences in the ease of syntactic processing.

The strength of harmonic series predicts the degree to which native German speakers are sensitive to word order violations, i.e., the grammaticality manipulation we selected for our experiment. The online comprehension of German relies substantially on word order within and across phrases [37]. However, within the Germanic language family, English, sensibly, assigns word order more weight for comprehension than German does [38]. We therefore predict that the sensitivity to word order violations, as reflected by the strength of neural harmonics, should be larger for languages which strongly depend on word order for comprehension, such as English, and smaller for morphologically richer languages, such as Italian, in which comprehension is locally bound by rules of declension. This perspective opens new, exciting avenues of cross-linguistic testing of different types of grammaticality constraints in multilanguage acquisition settings, as well as for language recovery in clinical settings.

By correctly characterizing the correspondence between linguistic functions and brain data, we contribute to linguistic theory by proving no evidence for the existence of a superordinate phrasal category - *sPhrase* - cycling through all phrases regardless of their lexical head can be posited. A proposal along these lines was first put forward by Chomsky [30], following the observation of similarities in the internal structure of NPs and VPs (see also [39–40]). In this respect, a frequency tagging approach to abstract multi-word categories has the potential to contribute directly to linguistic theories. However, there are also limitations to our experimental setup and analysis. First, FFT-base analyses of repeated sentence structures convey information on the *result* of top-down comprehension processes - that is, whether syntactic representations have been accessed from long-term memory -, but cannot provide any evidence for the incremental bottom-up processes combining words into phrases, and phrases into sentences (for a different take, see [23]). More generally, endogenous rhythms can in principle simply reflect the outcome of a decoding process that originates when participants encounter the first few trials, and is just top-down replicated for the rest of the trials. Second, endogenous neural rhythms generated via frequency tagging are uninformative relative to the temporal dynamics of internal representations in continuous speech perception. Third, the distinction between linear speech perception and hierarchical language comprehension sits well with our stimuli, as we use of monosyllabic and simple disyllabic units, but must not be overstated, as in many languages morphological processes contributing to word formation also have a hierarchical nature.

The delta band (0.5-4 Hz) and infra-slow (< 0.5 Hz) rhythms we measured lie within natural spectral niches for linguistic operations because their extended time windows match the ranges of spontaneous language segment duration [41–43]. However, delta rhythms also robustly entrain to salient events in continuous speech [44–45], as well as to meaningless word lists [22]. Frequency tagging offers a way to partition the contribution of language knowledge from that of speech, and is easily applicable to the study of top-down knowledge effects in other domains, such as music perception. An interesting point for future research pertains to the type of top-down knowledge that frequency tagging can tap into, from capturing complex syntactic structures (combining sentences within sentences) to detecting degrees of rule violations as a function of neural response strength.

In conclusion, we have shown that internally constructed neural rhythms and their harmonic series reflect the top-down application of syntactic rules determining the temporal boundaries of phrases and sentences, independently of entrainment to word rate.

## Materials and methods

### Participants

Thirty-two young adults participated in the Experiment (Age range: 19-30, 8 males). Sample size was determined using values derived from the inspection of Figure 1c in Ding et al. [20]: N = 16, mean response across peaks ± 12 dBs (SD ± 8 dBs), effects size = 1.5, power = 0.99. We hypothesized a reduction of 50% in effect size due to EEG volume conduction, leading to a required sample size of 30 with alpha = 0.05. The EEG data from one participant were not faithfully recorded, and were thus discarded. All analyses were run on the remaining participants (N = 31): they were all German native speakers, right-handed (Oldfield Test), and self-reported normal hearing, normal-to-corrected vision, no medical history of treatments affecting the central nervous system, and no psychiatric disturbances. They were compensated with 10 euros per hour (~3 hours, including instruction, EEG montage and cleaning). All experimental procedures were approved by the Ethics Committee of the Max Planck Society, and were undertaken with a written informed consent signed by each participant.

### Stimuli

To reduce implicit word- and phrase-level prosodic cues, word stimuli were individually generated using the MacinTalk Text-to-Speech Synthesizer (OSX El Capitan, voice Anna, f0 set at 220 Hz), and then pitch-flattened to 220 Hz across all time points (fundamental frequency estimated using an autocorrelation approach, Praat, v. 6.0.16, www.praat.org). If an original stimulus was longer than the relative SOA, its duration was compressed to fit the stimulation window while preserving pitch.

Individual trials were composed of ten sentences, identical from the viewpoint of main phrasal units, but differing in the types of lexical heads and internal composition of phrases. Sentences were randomly selected from six predefined lists of 50 sentences, one list per sentence structure (see Procedure, and Supporting Information). Within a trial, no sentence was ever repeated. There were two types of trial: regular trials, which contained only grammatically correct sentences; and irregular trials, which contained also two ungrammatical sentences, obtained by randomly shuffling word positions across two successive sentences. There were fifty trials per sentence structure, half were regular and half irregular, thereby yielding unbiased estimates of accuracy in discriminating regular from irregular trials, a task akin to classic grammaticality judgments.

### Procedure

Participants sat comfortably in an IAC 40a sound attenuating and electrically shielded recording booth (IAC Acoustics, iacacoustics.com), approximately 1m from an LCD computer screen. They were instructed to fixate a cross at the centre of the screen and listen attentively to each trial; When a trial ended, they were asked to determine whether it was regular or irregular. They were asked to be as accurate as possible, with no time limit. Participants used their right hand to press on two pre-defined buttons (arrow up regular, arrow down irregular) on a keyboard. Stimuli were delivered diotically at 75 dBs SPL via loudspeakers positioned at circa 1.2 m from participants, 25 cm to the left and right of the LCD screen. On presentation, stimuli were further attenuated using a fixed −20 dB SPL step, resulting in a comfortable and perceptually controlled environment. Brief rest periods between blocks were self-determined by each participant.

Trial presentation was blocked by sentence structure. Stimuli within a trial were presented at a constant Stimulus Onset Asynchrony (isochronous SOA): 250 ms for monosyllabic words, 500 ms for disyllabic words. There was no gap between sentences within a trial: Participants listened to a continuous flow of words equally spaced in time, further reducing prosodic cues at phrasal and sentence level. The main experimental manipulation pertained to the size - in number of words - of Noun Phrases (NP) and Verb Phrases (VP) composing each sentence structure. An NP is a phrase with a noun as head (e.g., “The girls”), while a VP has a verb as head (e.g., “play rugby”). NPs usually perform the grammatical functions of verb subject or verb object. In our experiments, all grammatical sentences contained an NP functioning as verb subject, and a VP, which either included a second NP functioning as a verb object or a different phrasal component. The Appendix lists all token sentences. See Figure 2 for an exemplary analysis of a sentence used in each condition. Stimulus sequences were created using custom scripts written in Matlab (R2015b, 64 bit, mathworks.com). Sequence delivery was controlled by Psychophysics Toolbox Version 3 (PTB-3, psychtoolbox.org) for Matlab, running on a Windows 7 computer (ASIO sound card for optimal stimulus latency control).

### Data Recording and Analysis

Behavioural accuracy data were subject to a signal detection theory approach, obtaining measures of task sensitivity (d’ = zscore(Hits) minus zscore(False Alarms)) and response bias (criterion = - 0.5*(zscore(Hits)+zscore(False Alarms))) for each participant and condition [32]. We used an actiCAP 64-channel, active electrode set (10-10 system, Brain Vision Recorder, Brain Products, brainproducts.com) to record electroencephalographic (EEG) activity at a sampling rate of 1KHz, with a 0.1 Hz online filter (12 dB/octave roll-off). Additionally, electrocardiographic (ECG) and electrooculographic (EOG) signals were recorded using a standard bipolar montage. All impedances were kept below 10 kOhm. Data were recorded with an on-line reference to FCz channel, and offline re-referenced to the average activity of all scalp channels, and downsampled to 250 Hz. Using the EEGLAB toolbox for Matlab [46] (sccn.ucsd.edu), continuous EEG data were first visually inspected to remove large non-stereotypical artifacts (e.g., sudden head movements, chewing), digitally filtered at 35 Hz low-pass (Kaiser window, beta = 5.65326, filter order 93, transition bandwidth 10 Hz), high-pass filtered at 1 Hz (filter order 455, transition bandwidth 2 Hz) to ensure data stationarity, submitted to an automatic artifact rejection based on spectrum thresholding (threshold = 10 standard deviations, 1-35 Hz), and then decomposed into Independent spatial components using the Infomax algorithm, which allows for optimal source separation [47]. The resulting Independent Components (ICs) were tested using the SASICA toolbox for EEGLAB [48]: ICs reflecting blinks/vertical eye movements, lateral eye movements and heart-beat, were detected by means of a correlation threshold (0.7) with bipolar Vertical, Horizontal EOG, and ECG channels, respectively, and found to be present in all participants (range: 1-3 ICs for vertical eye movements, 1-2 ICs for horizontal, 1 IC for heart beat). To reduce inter-trial signal variability, ICs reflecting muscle artifacts were identified via autocorrelation (threshold = 0, lag = 20 s), while ICs reflecting focal topography was used for bad electrodes (= 7 standard deviations threshold relative to the mean across electrodes). ICA results were copied back to the original EEG datasets (0.1 Hz high-pass), and ICs marked as artefactual were rejected before trial epoching. Each ten-sentence trial varied in duration between 10 seconds (4 words per sentence, running at 4 Hz) and 25 seconds (5 words per sentence, running at 2 Hz). Individual trial power estimates were extracted for each electrode from Hann-windowed and standardized data using the complex modulus of the Fast Fourier Transform, correcting for the Hann window loss of power (sqrt(1.5)). FFT trials were then transformed into phase coherence estimates by dividing by the amplitude of the signal, summating the angles, taking the absolute value and normalizing by the number of trials for each sentence type and participant. Signal-to-noise ratio estimates (SNR) were calculated for each frequency point by normalizing its value by the average of two samples before and two samples after it.

To verify the presence of significant neural peak and the effect of syntactic manipulations, a series of T-tests, repeated measures analysis of variance (rmANOVAs) and Bayesian rmANOVAs [49], were run, verifying the evidence for the categorical harmonics model. False-discovery-rate (FDR) correction with Q value = 0.05 was applied in all cases of multiple comparisons [50]. A machine learning approach to trial classification was run on the standardized (z-scored) phase coherence scores for each participant and each harmonic series peak, with the addition of the word rate peak. Three independent, numerically balanced subsets (“folds”) were created for each peak from the scores of all participants (“samples”), and an iterative process was repeated independently for each fold, while the other two were used a training set to train an LDA (Linear Discriminant Analysis) classifier [51]. Electrodes counted as features in this analysis. The resulting classifiers were tested by predicting whether a trial label was irregular or regular. Accuracy was quantified using the area under the curve of the receiver-operating characteristic (ROC AUC)

This analysis was repeated three times, with each trial subset serving as testing set once. A Montecarlo resampling approach (1000 repetitions) was used to obtain 95% confidence intervals, by shuffling the order of the trial labels and replicating the LDA classification.

## Data Availability

Anonymised pre-processed datasets for this experiment, as well as the relevant analysis scripts, are in the process of being made available in OSF (https://osf.io/rqfws/, currently private).

## Acknowledgements

The authors would like to thank Cornelius Abel, Jana Gessert, Freya Materne, Claudia Lehr, Alexander Lindau, Georg-Friederich Paasch for help with stimulus creation and data collection, and Dobromir Dotov for critical feedback on data analysis.

